# Honey Bees Find the Shortest Path: A Collective Flow-Mediated Approach

**DOI:** 10.1101/2022.06.27.497822

**Authors:** Dieu My T Nguyen, Golnar Gharooni Fard, Michael Iuzzolino, Orit Peleg

**Affiliations:** University of Colorado Boulder

**Keywords:** Honey bees, Swarm intelligence, Self-organization, Problem solving, Traveling salesman problem

## Abstract

Honey bees (*Apis mellifera* L.) are social insects that makes frequent use of volatile pheromone signals to collectively navigate unpredictable and unknown environments. Ants have been shown to effectively use pheromone trails to find the shortest path between two points, the nest and the food source. The ant pheromone trails are accomplished by depositing of pheromones which are then diffused passively, creating isotropic (i.e., non-directional and axi-symmetric) signals. In this study, we report the first instance of the honey bees’ ability to solve the shortest path problem to localize the queen and aggregate around her by using a collective flow-mediated scenting strategy. In this strategy, individual bees not only emit pheromones, but also fan their wings to actively direct the flow of the signals, providing colony members with directional messages to the queen’s location. We use computer vision and deep learning approaches to perform automatic and accurate image analysis. As a result, we quantify the number of bees in the short and long paths, and show that the short path is frequented by significantly more bees over time. We also reconstruct attractive surfaces using the positions and directions of scenting bees, and show that this surface is more “attractive” along the short path and around the queen as scenting bees send out directional messages and the swarm makes their way to the queen. Overall, we show that the honey bees can effectively use the collective scenting behavior to overcome local and volatile pheromone communication and find the shortest path to the queen.

## 1 Introduction

In social insect colonies, members must effectively communicate to navigate unpredictable and unknown environments. Pheromones, or volatile chemical messages, are one of the prevalent signals that insects employ to exchange information and coordinate group processes, such as foraging or aggregating [1, 7, 15]. Although prevalent, pheromones decay rapidly in time and space, limiting the range and duration of information exchange. Thus, social insects must collectively solve this problem by creating effective and robust communication networks.

Ant pheromone trails are a well-known example of a collective, self-organized solution to the traveling salesman problem of finding the shortest route between the nest and a food source [3, 13]. Ants employ the method of passively depositing pheromones in their trajectories, producing isotropic (i.e., non-directional and axi-symmetric) signals. These chemical signals are reinforced on a given path over time as ants traverse the path in both directions, leading to the collective choice of the shortcut. Additionally, previous studies have shown that, a true slime mold, the plasmodium of *Physarum polycephalum*, can find the shortest path through a maze or connect different arrays of food sources in an efficient manner with low total length [9]. Similar to the ants, these single-celled amoeboid organisms employ passive signaling without centralized control or global information to also form networks with the efficiency, fault tolerance, and cost comparable to those of the real Tokyo rail system [14].

In this work, we aim to show that another species employing chemical signaling is capable of solving the shortest route problem: the honey bees (*Apis mellifera* L.). In our previous study of how honey bees can search for and swarm around the queen, we show that beyond passive depositing of chemical signals seen in the ants and amoeba, honey bees organize into a communication network and “scent” to emit pheromones and fan their wings to actively direct the signal flow and expanding the reach of the signals, providing colony members with directional messages of the queen’s location [10]. Now, we investigate whether the honey bees can solve the shortest path problem in the swarming context using the directional pheromone signaling strategy. To that end, we present worker bees with a Y-maze consisting of a short and a long path to the queen, who is caged and placed in a fixed location at the opposite end of the maze. This maze is inspired by the design in the classic work on the self-organized shortcut in ants above-mentioned [3]. We also ran a control experiment by presenting the bees with a mirrored version of the maze, to show the reproducibility of the behavior and eliminated any directionality bias of the bees’ behavior. We then track the search behavior of individuals and the swarming behavior of the group over time with computer vision and machine learning methods to automatically and accurately detect the bees’ location and their directional scenting behavior. Through these experiments, we aim to assess whether and how the honey bees employ a collective flow-mediated approach with volatile signals to solve the problem of finding the shortest path to a target, the queen, for aggregating.

## 2. Methods

### 2.1 Experimental setup

Honey bees have been shown to scent while standing stationary on a surface [8]. Thus, our experimental arena (50 cm × 50 cm × 1.5 cm) on top of a backlight board is semi-two-dimensional to prevent flying for easier handling and analysis. Wood blocks are used to build a maze with a short and long path (35 and 65 cm, respectively). See Fig. 1A for an image of the experimental setup, in which the bees are recorded from an aerial view with a video camera (4k resolution, 30 fps). Before the experiments, the queen and worker bees are collected from a colony. Workers are isolated from their original queen for 24 hours, then introduced to a different queen in a cage (10.5 × 2.2 × 2.2cm) from our queen bank and stayed with her for another 24 hours. In each experiment, the caged queen is placed at the right end of the maze consisted of two paths. Worker bees are placed at the other end. A sheet of plexiglass is placed on top of the arena to enclose the space. We regularly monitor temperature to sure the heat from the backlight board does not exceed 32 − 35°C and does not affect the fanning behavior. We perform three trials with the maze orientation shown in Fig. 1A, and three more trials with a mirrored version of this maze in which the short path is at the top and the long path is at the bottom. The ratio of the long and short paths is approximately 2, consistently with the ratio used in the maze in the classic ant paper [3]. We use different sets of bees for each trial so that our replications of the maze do not increase the capacity of the colony to find the preferred path (i.e., to eliminate learning). We also wash the cover with soap and water and use Lysol spray for the arena and maze in between trials to eliminate any leftover pheromone from the previous trial.

**Fig. 1.**
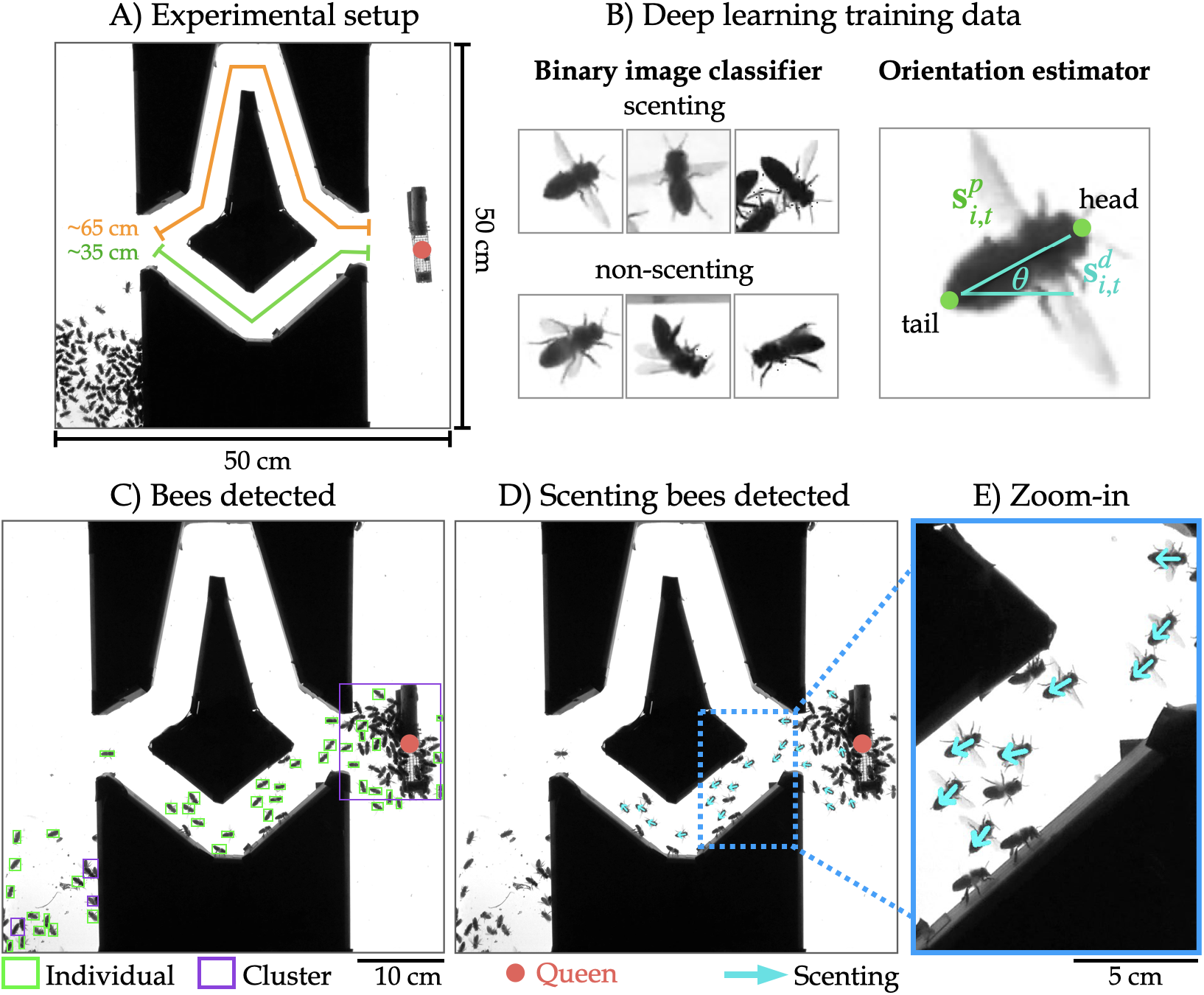
Experimental setup and image analysis with machine learning. A) Experimental setup of the semi-twodimensional area of 50 × 50 × 1.5 cm, in which the short path (green) is approximately 35 cm in length and the long path (orange) is approximately 65 cm. B) Examples of training data for two convolutional neural network models, the binary image classifier to classify scenting and non-scenting bees and the orientation estimator. C) Example detection of individual bees (green boxes) and clusters (purple boxes). D) Example detection of scenting bees and their scenting directions (teal arrows pointing from head to tail). E) Zoomed-in part showing example annotated scenting bees with wide wings.

### 2.2 Bee detection & scenting recognition

To automatically detect scenting bees and their scenting directions in the videos, we employ computer vision and deep learning approaches originally presented in detail in [10]. First, to detect individual bees, we extract images at 1 fps and use Otsu’s method to adaptively threshold the images [11, 12]. Morphological transformations (opening) are iteratively applied to remove noise and separate individual bees from clusters of bees that are difficult to parse out [2]. The connected component algorithm is then applied to obtain the components’ centroids (x, y positions) and areas. Large clusters are filtered out by area to isolate individual bees. See Fig. 1C for an example image annotated with individuals (green boxes) and clusters (purple boxes).

Second, to binarily classify individual bees as scenting or non-scenting, we train a ResNet-18 convolutional neural network (CNN) model [6] using 28,458 labeled images [10]. Examples of scenting and non-scenting bees are shown in Fig. 1B under “Binary image classifier.” The model is trained with augmented data (horizontal and vertical flipping, brightness adjustments, scaling, translation, and rotation) and balanced sampling to combat the class imbalance (9:1 non-scenting to scenting) for 1203 epochs with early stopping to prevent overfitting. On the test set, the model achieves 95.17% accuracy, indicating that our model can generalize to unseen data.

Lastly, we use the same ResNet-18 CNN for orientation prediction to provide us with the scenting directions. While the scenting classifier model outputs binary categorical labels, the loss function here is modified for the model to output continuous values for the predicted angles: 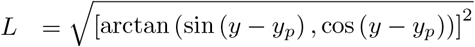, where *y* is the true label, and *y*_*p*_ is the model’s prediction. We created a labeled dataset of 15,435 images, each image with head and tail positions from which we compute the ground-truth orientation angle [10] (Fig. 1B under “Orientation estimator”). On the test set, this model achieves 96.71% with 15*°* of error tolerance. Fig. 1D and E show the complete result of detecting individual bees, detecting scenting bees out of the individual bees, and estimating the scenting directions (teal arrows).

### 2.3 Time-series analysis

From the computer vision pipeline above-described, we have the position and scenting information of the bees to extract time-series data. Per frame, we obtain the number of scenting, non-scenting, and all bees in the short and long path. In the results, we present the cumulative sum of the three types of bees as a rolling average with a window size of 60 frames or seconds.

### 2.4 Attractive surface reconstruction

To correlate the scenting events with the spatiotemporal density of bees, we reconstruct an attractive surface for each frame of a video, with a method originally presented in detail in [10]. To describe briefly, for each scenting bee *i* at time *t*, we define its position as 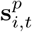, and its direction of scenting as 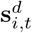 (unit vector). Assuming the scenting bees provide directional information to non-scenting bees, we treat 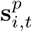 and 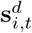 as a set of gradients that define a minimal surface of height *f* (*x, y, t*). Thus, *f* (*x, y, t*) corresponds to the probability that a randomly moving non-scenting bee will end up at position (*x, y*) by following the scenting directions of scenting bees:

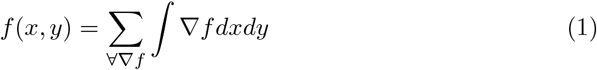

where 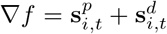. We regularize the least squares solution of the surface reconstruction from its gradient field, using Tikhonov regularization [4, 5].

## 3 Results

To characterize how the honey bees collectively behave when presented with two paths to the queen, we observe the experiments qualitatively and quantitatively over time. In Fig. 2, we show an example experiment with snapshots of the bees interacting in the semi-twodimensional arena with a caged queen. As shown in Fig. 2A, the worker bees perform early exploration of the two paths (t=455 sec), then converge and scent on the short path to signal the queen’s location to the rest of the swarm (t=570 sec). By approximate 820 sec, most of the bees have aggregated around the queen. In Fig. 2B, we present the corresponding attractive surfaces *f* reconstructed from the scenting bees’ positions and directions. The strongest signals tend to lie along the short path and around the queen.

**Fig. 2.**
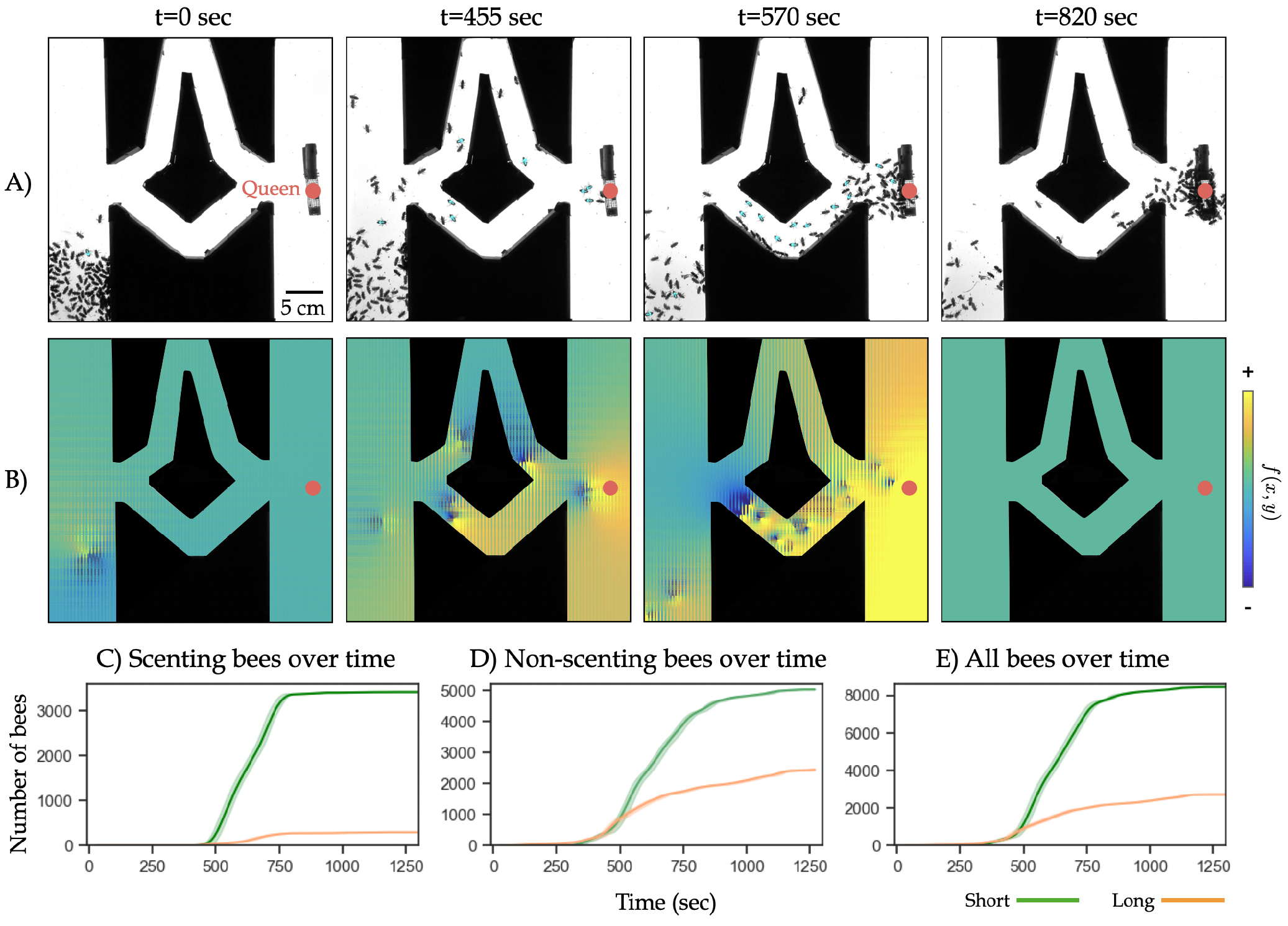
Time series of an example maze experiment. A) Snapshots of an example experiment in a semi-two-dimensional arena where worker bees are placed at one end of the maze with short and long paths, and the queen is located at the opposite end. Over time, the bees communicate via the collective scenting behavior to find the short path and swarm at the queen. B) Snapshots of the corresponding surfaces *f* according to Eq. 1 to show how the scenting events correlate to the spatial-temporal density of the bees. C) The cumulative sum of the number of scenting bees on the short (green) and long (orange) paths over time, shown as a rolling average with window size of 60 seconds. D) The cumulative sum of the number of non-scenting bees on the short (green) and long (orange) paths over time, shown as a rolling average with window size of 60 seconds. E) The cumulative sum of the number of scenting and non-scenting (i.e. all) bees on the short (green) and long (orange) paths over time, shown as a rolling average with window size of 60 seconds.

We also quantify the temporal dynamics of the search for the shortest path and aggregation around the queen. In Fig. 2C, we show the cumulative sum of the number of scenting bees on the short (green curve) and long (orange) paths over time, shown as a rolling average with window size of 60 seconds. In Fig. 2D and E, we show the data for non-scenting bees and all bees (e.g. scenting and non-scenting), respectively. There are significantly more bees of all kinds on the short path, suggesting that the collective scenting communicates the preferred path to the queen to the rest of the swarm.

We perform six total trials of this experiment and show the time-series data for scenting bees and all bees in Fig. 3 for all trials. In three trials (1 to 3), we present the bees with the maze as shown in the example in Fig. 2. In the rest of the trials (4 to 6), we flip the maze to prevent any memory or directional bias. In trials 1 to 5, the number of scenting and all bees on the short path are consistently higher than on the long path. In the outlier trial 6, we observe the opposite dynamics. Note that we record the number of bees on the two paths per video frame. In trial 6, the cumulative sum may include bees that travel the paths back and forth rather than most bees forming a swarm at the queen’s location as seen in trials 1 to 5.

**Fig. 3.**
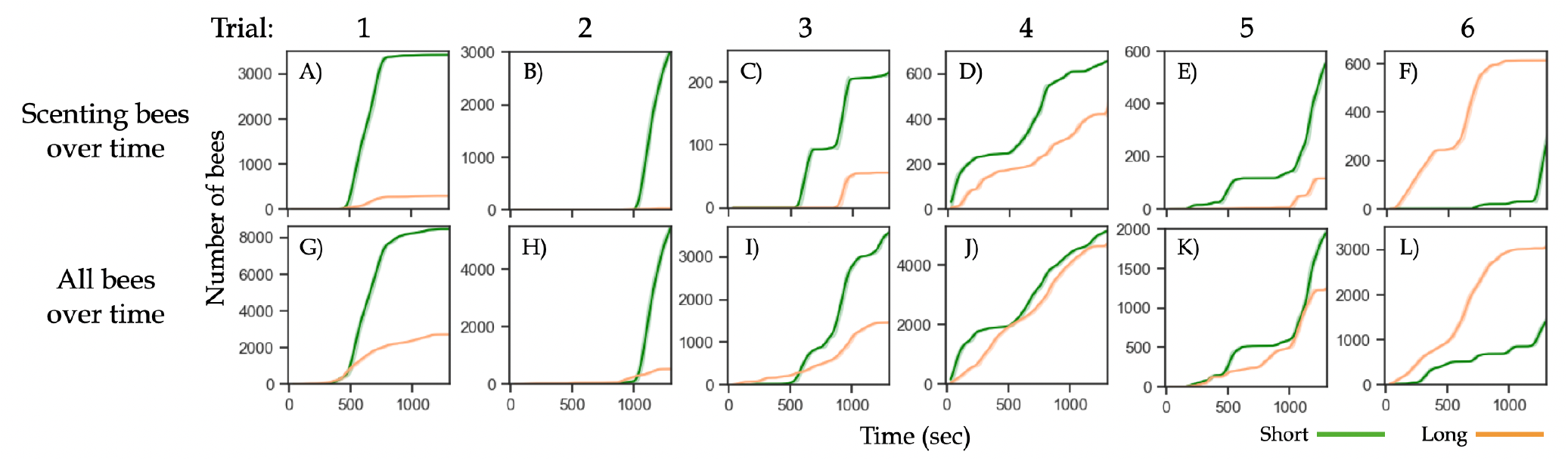
Time series data of all six trials. Trials 1 to 3 consist of a maze with the orientation presented in Fig. 2. Trials 4 to 6 consist of a mirrored maze in which the short path is at the top and long path is at the bottom. A–F) The cumulative sum of the number of scenting bees on the short (green) and long (orange) paths over time, shown as rolling average with window size of 60 seconds, for all six trials. Trial 1 is the example shown in Fig. 2. Besides in trial 6, there are more scenting bees on the short path than on the long path. G–L) The cumulative sum of the number of all bees (e.g. scenting and non-scenting) on the short (green) and long (orange) paths over time, shown as rolling average with window size of 60 seconds, for all six trials. Trial 1 is the example shown in Fig. 2. Besides trial 6, there are more bees overall on the short path than on the long path.

## 4 Discussion & Conclusion

In this work, we investigate how the honey bees use a collective flow-mediated communication approach to find the shortest path to the queen, the target for swarm formation. After some initial exploration stage of the environment, the bees can collectively converge on the short path to cross and move towards the queen’s location. We observe the active chemical signaling behavior of scenting in the bees, especially on the short path as the animals emit and propagate the information to swarm members. As a result, the short path to the queen is often populated by more bees and becomes more “attractive” as the swarm members effectively communicate the collective preference for the more optimal path to the queen.

In our experiments, we keep the ratio of the long and short path to an approximate value of 2, which has been shown in the ants to be a value where the collective choice of the shortcut is significant and consistent, a result we also observe in the bees. We note that the original work on the ants also experimented with ratio values of 1 (i.e. the two paths are the same length) and an intermediate value of 1.4. When the ratio is 1, the choice between two paths is roughly 50%. At the ratio of 1.4 and above, the ants show a clear preference for the shortest path. To extend our experiments with the bees, we aim to construct and test mazes with paths of different ratios to set a control (with a ratio of 1) and to explore the ratio threshold at which we can observe the bees successfully finding the shortest path. This may also be investigated in modifying the agent-based model of the collective scenting behavior in bees for swarming presented in [10]. The model allows exploration of how behavior parameters, such as the pheromone detection threshold and the magnitude of the directional signals, change with a more complex environment in order for the bees to adapt and successfully aggregate.

Altogether, we present the first findings of the honey bees’ capability of finding the shortest path. While individual bees are only equipped with local navigational information and spatiotemporally limited communication signals, they interact with one another via the reliable scenting strategy that harnesses the power of the collective.

## Acknowledgements

This work was supported by the National Science Foundation Graduate Research Fellowship (NSF GRFP) under Grant No. DGE 1650115 (D.M.T.N.) and Physics of Living Systems Grant No. 2014212 (O.P.). Any opinion, findings, and conclusions or recommendations expressed in this material are those of the authors(s) and do not reflect the views of the NSF. We also acknowledge the BioFrontiers Institute (internal funds), the Interdisciplinary Research Theme on Autonomous Systems (O.P.).

## Notes

### Competing Interest Statement

The authors have declared no competing interest.

## References

1. Conte, Y.L., Hefetz: Primer pheromones in social hymenoptera. Annu. Rev. Entomol. 53(1), 523–542 (2008)

2. Dougherty, E.R.: An introduction to morphological image processing. Society of Photo Optical (1992)

3. Goss, S., Aron, S., Deneubourg, J.L., Pasteels, J.M.: Self-organized shortcuts in the argentine ant. Naturwissenschaften 76(12), 579–581 (1989)

4. Harker, M., O’Leary, P.: Least squares surface reconstruction from measured gradient fields. CVPR pp. 1–7 (2008)

5. Harker, M., O’Leary, P.: Least squares surface reconstruction from gradients - Direct algebraic methods with spectral, Tikhonov, and constrained regularization. CVPR (2011)

6. He, K., 0006, X.Z., Ren, S., 0001, J.S.: Deep Residual Learning for Image Recognition. CVPR (2016)

7. Lensky, Y., Cassier, P.: The alarm pheromones of queen and worker honey bees. Bee world 76(3), 119–129 (1995)

8. McIndoo, N.E.: The Scent-Producing Organ of the Honey Bee 66(2) (1914)

9. Nakagaki, T., Yamada, H., Toth, A.: Path finding by tube morphogenesis in an amoeboid organism. Biophysical chemistry 92(1-2), 47–52 (2001)

10. Nguyen, D.M.T., Iuzzolino, M.L., Mankel, A., Bozek, K., Stephens, G.J., Peleg, O.: Flow-mediated olfactory communication in honeybee swarms. Proceedings of the National Academy of Sciences 118(13) (2021)

11. Otsu, N.: A threshold selection method from gray-level histograms. IEEE transactions on systems, man, and cybernetics 9(1), 62–66 (1979)

12. Sezgin, M., Sankur, B.: Survey over image thresholding techniques and quantitative performance evaluation. Journal of Electronic imaging 13(1), 146–165 (2004)

13. Shah, S., Bhaya, A., Kothari, R., Chandra, S., et al.: Ants find the shortest path: a mathematical proof. Swarm Intelligence 7(1), 43–62 (2013)

14. Tero, A., Takagi, S., Saigusa, T., Ito, K., Bebber, D.P., Fricker, M.D., Yumiki, K., Kobayashi, R., Nakagaki, T.: Rules for Biologically Inspired Adaptive Network Design. Science 327(5964), 439–442 (2010)

15. Trhlin, M., Rajchard, J., et al.: Chemical communication in the honeybee (apis mellifera l.): a review. Vet. Med 56(6), 265–73 (2011)

